# Evidence of parental care as a newly identified reproductive isolating barrier

**DOI:** 10.1101/2025.04.18.649590

**Authors:** Colby Behrens, Meghan F. Maciejewski, Alexandra Sumarli, Gina Lucas, German Lagunas-Robles, Kieran Samuk, Alison M. Bell

## Abstract

Variation in behavior can contribute to reproductive isolation by preventing gene flow among populations. Here, we tested the novel hypothesis that parental care, when dysregulated, can function as a reproductive isolating mechanism in three-spined stickleback fish (*Gasterosteus aculeatus*). In the typical “common” stickleback ecotype, males provide care to their offspring by fanning with their pectoral fins and defending their nest. In contrast, a divergent “white” stickleback ecotype has evolutionarily lost care and disperses embryos into the surrounding environment. We examined how paternal care from common, white, and F1 hybrid fathers influenced offspring survival. We detected no intrinsic incompatibilities in embryos, but F1 hybrid fathers exhibited dysregulated parental care that coincided with decreased survival of parented offspring. The increased offspring mortality may be explained by further dysregulation of feeding and parenting circuitry, as F2 hybrids exhibited significantly higher rates of post-fertilization filial cannibalism than common or white fathers. Additionally, despite strong divergence in nesting-building and courtship behavior, F1 hybrids achieved mating success at a similar rate as male commons and whites, suggesting that prezygotic barriers against hybrids may be weak. The observed postzygotic isolation, and potentially weak prezygotic isolation in lab-based studies, suggest that hybridization is likely occurring at low rates in the wild. Population genetic analysis supported this, as low proportions of putative hybrids were detected in sympatric sites. Together, these results may help explain why genetic divergence between these phenotypically distinct ecotypes is low and provide evidence that dysregulated parental behavior can act as a newly discovered postzygotic reproductive isolating barrier.

## INTRODUCTION

Behavior can directly affect survival and reproductive success and therefore plays an important role in evolutionary processes, including the evolution of reproductive isolating barriers (Coyne and Orr 2004). For example, behaviors like courtship displays, territoriality, and song can facilitate speciation if they are involved in preventing the formation of zygotes between different populations, thereby acting as prezygotic reproductive isolating barriers (Silberglied and Taylor 1978; Coyne and Orr 1989; Fitzpatrick and Gray 2001; Lackey and Boughman 2017). Behavior can also act as a postzygotic isolating barrier if, for example, hybrids display altered behaviors that are important for survival or reproductive success, such as intermediate migration routes (Helbig 1991; Delmore and Irwin 2014), hampered cognition (McQuillan et al. 2018) or dysregulated courtship behavior that falls outside of the normal adaptive range (Pashley and Martin 1987; Noor 1997; Gottsberger and Mayer 2007; Clark et al. 2010).

Here, we investigate dysregulated parental care as a previously undescribed behavioral postzygotic isolating barrier in animals. Although parental care is relatively rare in nature, it has evolved independently multiple times and is critical for the development and survival of offspring in those groups (Clutton-Brock 1991). Successful rearing often requires proper expression of a suite of behaviors, including resources allocation, brood hygiene, and predator defense, and disruption of any of these could reduce offspring survival. Importantly, rearing failure could also result from the exaggeration of behaviors. For example, though filial cannibalism is relatively common in nature (Polis 1981; Klug and Bonsall 2007), a moderate increase in this behavior could significantly increase mortality in a clutch. Therefore, hybrids that exhibit altered or dysregulated parental behavior in any number of these traits may have low fitness, even if no gametic or other intrinsic incompatibilities are present.

In this study, we test the hypothesis that dysregulated parental behavior can act as a postzygotic reproductive isolating barrier in three-spined stickleback fish (*Gasterosteus aculeatus*), a classic model for speciation (McKinnon and Rundle 2002). Stickleback are famous for their propensity to adapt to new environments, and the repeated colonization of freshwater habitats by marine ancestors has resulted in the radiation of sticklebacks throughout the Northern hemisphere. Studies of “species pairs” of phenotypically divergent populations, e.g., benthic-limnetic (McPhail 1992; Arnegard et al. 2014; Keagy et al. 2016; Bay et al. 2017), lake-stream (Hendry et al. 2002; Räsänen and Hendry 2014), freshwater-anadromous, (Hay and McPhail 1975; Ziuganov 1995; Aguirre et al. 2022) and Japan Sea stickleback (Kitano et al. 2007; Kitano et al. 2009) have offered strong support for the importance of ecological drivers of divergence in sticklebacks.

A less-studied pair of populations are the “white” and “common” stickleback ecotypes found in Nova Scotia, Canada. Whites and commons occupy the same trophic niche (Samuk 2016), yet they exhibit markedly divergent reproductive behavior, especially parental care. As is typical of the family Gasterosteidae, males of the common stickleback ecotype provide parental care for several days by aerating their embryos via pectoral fanning and by removing decaying eggs, which is critical to preventing fungal infections (Wootton 1976). In contrast, the divergent white ecotype has evolutionarily lost care. After fertilizing the eggs, white stickleback fathers remove embryos from the nest and disperse them into the surrounding environment (Blouw and Hagen 1990; Blouw 1996; Behrens et al. 2024). Male whites and commons differ notably in several other traits important for reproduction including nest architecture (male whites build relatively loose nests) and courtship behavior (male whites engage in high rates of female-directed courtship behavior (Jamieson et al. 1992; Haley et al. 2019; Behrens et al. 2024)). Female whites and commons have also diverged in reproductive traits, with female whites producing relatively large clutches of small eggs (Grant 1993; Behrens et al. 2025). Female whites produce clutches of eggs that do not stick together (Grant 1993), which may facilitate the embryo scattering behavior of male whites and prevent the spread of fungus throughout the entire clutch.

Population genetic analyses suggest that the ecotypes diverged recently (<12K years) and are genetically distinct, but their genome-wide differentiation is low (F_ST_ ∼0.02) and there is likely gene flow between them (Samuk 2016; Sumarli, A. & Samuk, K. personal communication). The populations frequently occur in sympatry but there are currently no known sites where only whites are found. There is no evidence of altered embryonic development or inviable gametes in F1 hybrids or their offspring: laboratory-generated F1 and F2 hybrids are fertile and produce viable offspring (Blouw 1996; Behrens et al. 2024; Behrens et al. 2025), suggesting that hybridization could be occurring in the wild. It is possible that partially non-overlapping breeding seasons and/or assortative mating contribute to prezygotic isolation in this system, but evidence is mixed (Blouw and Hagen 1990; Corney and Weir 2023). Altogether, it is unclear what processes generated and maintained such extensive phenotypic divergence without a similar genetic divergence, and the rate at which hybrids occur in natural populations is unknown.

Here, we test the novel hypothesis that dysregulated parental behavior can be a form of behavioral sterility (Coyne and Orr 2004) and act as an extrinsic postzygotic reproductive isolating barrier by preventing F1 hybrids from achieving reproductive success. In the white-common stickleback system, this predicts that even if the white and common ecotypes hybridize in the wild, the two ecotypes fail to mix completely because F1 hybrid males exhibit dysregulated paternal behavior that results in low offspring survival. This hypothesis is motivated by a previous study showing that F1 hybrid white x common males perform a mixture of typical white and common behaviors after mating, with relatively low levels of the behaviors characterizing each strategy, (i.e. low levels of dispersal behavior and low levels of fanning (Blouw 1996; Behrens et al. 2024)). Given the importance of paternal care for offspring survival in this species (Wootton 1976), we predicted that dysregulated behavior in F1 hybrid fathers would result in low offspring survival.

To test this hypothesis, we quantified multiple potential mechanisms of reproductive isolation in this system. First, we compared the survival of embryos reared by white or common fathers to the survival of embryos that received care from F1 hybrid fathers. If dysregulated paternal behavior acts as a postzygotic reproductive isolating barrier in this system, then we predicted that embryos that received care from F1 hybrid fathers would have lower survival than embryos that received care from white or common fathers. In addition to measuring offspring survival, we also measured paternal behavior to gain insights into the behavioral mechanisms that may contribute to behavioral sterility, including behaviors that reduce care or increase offspring mortality. To control for potential intrinsic barriers, we measured the survival of offspring that were reared artificially, which generates high rates of embryo survival if there are no genetic incompatibilities or gametic sterility (Day et al. 1994), and compared the survival of artificially-reared offspring sired by F1 hybrid fathers to the survival of offspring sired by white or common males.

Like many fishes, male stickleback often consume eggs from conspecifics’ nests (FitzGerald 1991) as well as their own (filial cannibalism). The causes of filial cannibalism in sticklebacks are not well understood (see Mehlis et al. 2009), but we suspected that if paternal care was dysregulated in hybrid fathers then it could manifest as higher than usual rates of filial cannibalism, as feeding and parenting neural circuits are closely linked in vertebrate brains (Fischer and O’Connell 2017). If hybrid males possess dysregulated parental circuits, this dysregulation could also elevate feeding-related behaviors like filial cannibalism. We therefore examined rates of filial cannibalism by recently mated male commons, whites, and hybrids, and predicted that hybrids would exhibit higher rates of cannibalism than either parental ecotype.

For completeness, we also explored the possibility of mate choice acting as a prezygotic barrier against hybrid males. White and common males employ unique courtship and nesting strategies throughout the reproduction season (Behrens et al. 2024), and the intermediate displays of hybrid males could appear unattractive to potential female mates. We therefore examined the mating success of hybrid males and investigated the potential role of nest architecture as an isolating barrier preventing the formation of backcross hybrids.

Finally, we examine the prevalence of hybrids across natural populations of white and common stickleback. Though gene flow is likely ongoing in this system (Samuk 2016), the presence and rate of hybridization has never been examined in wild populations and could inform the interpretation and implications of this lab-based study for isolating barriers in natural populations. Here, we bring evidence to bear from a population genomic survey which shows that hybridization occurs in the wild.

## METHODS

### Animal collection and husbandry

In 2023, three-spined stickleback were collected from two sites in Nova Scotia, Canada. White sticklebacks were collected at Salmon River (45°21’9.57’N 61°28’22.20’W) and common sticklebacks were collected from Blue’s Cove (45°53’55.7’N 61°05’11.1’W) using unbaited minnow traps. Because white stickleback are absent from Blue’s Cove, all males collected at that site were unambiguously classified as common stickleback. Both ecotypes occur at Salmon River, so we differentiated whites from commons using known differences in body size, nuptial coloration, and egg stickiness (Blouw and Hagen 1990; Behrens, Arredondo, et al. 2025). Hybrid stickleback are typically phenotypically intermediate (Behrens et al. 2024), and no morphologically intermediate individuals were collected at either site. Clutches of white, common, and a full reciprocal cross of F1 hybrids were then generated through artificial fertilization, where testes from sacrificed males were macerated and applied over clutches of eggs squeezed from females. Embryos were reared at the University of California Riverside (Methods S1) and later transported to the University of Illinois Urbana-Champaign for experimental testing.

### Sample collection for population genetic analyses

In June 2019, adult common stickleback were collected from Cherry Burton Road (N 46° 01.516’ W 64° 06.150’) and white stickleback were collected from Canal Lake (44°29’54.0”N 63°54’09.1”W) and transported to the University of Illinois in Urbana-Champaign before tissue collection. In the spring of 2022, random fin clips were obtained from 50 additional three-spined sticklebacks from each of four sites in Nova Scotia, two where white and commons are known to be sympatric (Canal Lake and Salmon River, coordinates above), and two where only commons are found (Blues Cove, discussed previous, and McIvor Road Junction, 45°56’09’N 61°04’06’W; Methods S2). There are no known sites where only whites are found.

### Behavioral phenotyping and offspring survival

Upon entering breeding condition, signified by red throats and blue eyes, 1-year-old male commons, whites, ♂C x ♀W F1s, and ♀C x ♂W F1s were housed in individual tanks and provided with nesting materials (Behrens et al. 2024). Male commons and whites were paired with gravid females from their respective populations, and males of both F1 hybrid cross types were mated to female commons or female whites, resulting in males in 6 different conditions, with n = 6 males per condition. Whites and commons (F0s) are expected to be more abundant than F1 hybrids in the wild, so we did not test intercross (F1 hybrid x F1 hybrid) conditions. We quantified the architecture of male nests if they were visible without manipulation. Briefly, nests received qualitative scores on four attributes (height, opening, sand content, location) that were combined in a single nesting metric (Behrens et al. 2024). We then introduced a live gravid female into the male’s tank to spawn via no-choice trials (Boughman 2001). The female was weighed, placed into a male tank, and removed if a mating did not occur within 15 minutes. If a mating did occur, the female was removed and reweighed to estimate the mass of the clutch. We tracked the number of females that were introduced to each male before successfully mating as a rough proxy for mate preference. After fertilization male parenting behaviors, including fanning and embryo dispersal (Wootton 1976; Blouw 1996; Behrens et al. 2024), were recorded twice daily for 15 minutes for four days with the program JWatcher (Blumstein et al. 2006) in order to gain insights into the behavioral determinants of successful reproduction.

Fathers were removed from their tanks four days after fertilization, which is when many parenting behaviors begin to peak (Behrens et al. 2024). This timepoint was chosen to accurately quantify embryo survival during peak parental care; therefore, differences in survival may be conservative estimates relative to effects observed in later stages of care. Males were euthanized and testes were removed to artificially fertilize an additional clutch of eggs that was artificially reared (Day et al. 1994); clutch size of artificially-reared clutches was manually counted immediately after fertilization. The same female ecotype was used for artificially- and father-reared clutches for each male.

Embryos from natural fertilizations were also collected at this time to estimate the survival of father-reared offspring. To collect embryos located in the nest, we placed the nest under a dissection microscope and sifted through algal strands with fine forceps. Care was taken to ensure that embryos were not dropped into the tank during nest removal. White and hybrid stickleback disperse embryos into the surrounding environment; therefore, a reliable estimate of offspring survival requires a survey of the entire tank. However, stickleback embryos are transparent and difficult to identify while resting on substrate, especially during early stages of development (Swarup 1958). To account for this, we removed tank substrate (e.g. gravel) in increments of <240 ml, spread the substrate loosely across small tanks (dimensions 7×4×5”), and used plastic dropper pipettes to push any embryos into the water column for detection and counting. Tanks from all males, including the non-dispersing commons, were surveyed. Healthy embryos are readily identifiable by eye capsules and blood circulation at this stage in development (Swarup 1958) so only living embryos were collected and saved. This method enabled estimation of embryo survival regardless of parenting strategy with high confidence. All embryos were then counted and artificially incubated until hatching.

### Filial cannibalism

To quantify rates of cannibalism, the stomachs of males were dissected and the number of eggs/embryos was counted within one hour after mating, when the critical decision about whether to disperse (whites) or care for embryos (commons) is made. Data on filial cannibalism by the common, white, or F1 hybrid males in the survival study was not available because their parenting behavior was observed for four days after mating, and rates of cannibalism are low at that point. A separate set of whites and commons (n=22 per ecotype) was assessed for filial cannibalism across 2021 and 2022 ,and filial cannibalism of F2 hybrids (originating from a single white ♀ x common ♂ cross, n=77) was quantified in 2021 (Behrens, Tucker, et al. 2025). We therefore test the hypothesis that hybridization influences filial cannibalism using filial cannibalism rates of F2 hybrids as an indirect proxy. Male whites and commons were paired with females from their own ecotypes and F2 hybrids were paired with female whites. Males from 2021 (common, white, and F2 hybrid) had body mass recorded immediately prior to dissection. Males were consistently fed in the morning prior to mating to prevent differences in satiation from affecting rates of cannibalism. The fertilization rate of cannibalized eggs was recorded to assess whether filial cannibalism was preferentially directed toward unfertilized eggs, which are susceptible to fungal infections that could threaten the entire clutch (Wootton 1976).

### Population genetics

#### DNA extractions & library prep

We performed standard phenol-chloroform DNA extractions (Sambrook et al. 1989) to extract DNA from the 200 randomly sampled individuals from 2022 (collected from Canal Lake, Salmon River, Blues Cove, and McIvor Road Junction) as well as 20 common sticklebacks (10 male, 10 female) from Cherry Burton Road and 20 white sticklebacks (10 male, 10 female) from Canal Lake collected in 2019.

DNA from the 200 randomly sampled individuals was normalized and prepared for double digest restriction-site associated DNA (ddRAD) sequencing using published methods (Brelsford et al. 2016), which implements elements proposed by Parchman et al. 2012 and Peterson et al. 2012, with the restriction enzymes PstI and MseI and removed small DNA fragments with Omega magnetic beads (OMEGA Bio-Tek). The remaining 40 samples were sent to the UC Davis DNA Technologies and Expression Analysis Core Facility for Whole Genome Sequencing (WGS) library prep. All sequencing (RAD and WGS libraries) was performed at the UC Davis DNA Technologies and Expression Analysis Core Facility on an Illumina High Seq-X. Variants were then called on RADseq and WGS reads (Methods S3)

#### Principal Component and Hybrid Analysis

The extremely low genetic differentiation between white and common sticklebacks precludes standard analyses aimed at hybrid classification (e.g. triangle plots). To identify potential hybrids, we performed principal component analysis (PCA) on the jointly filtered SNPs from the RADseq and WGS data. We used PCA instead of other commonly used model-based methods (e.g. ADMIXTURE) because it does not make any assumptions about a fixed number of discrete populations in our data. We first performed PCA on the total dataset. Because sex and ecotype identity were unknown in the randomly sampled fish, we assigned sex and ecotype genotypically. Sex was assigned by identifying individuals with high numbers of homozygous Y-linked SNPs in the non-PAR region of the Y-chromosome. Ecotype identity was assigned by examining the co-clustering of RADseq samples with WGS samples (which have known ecotype identity). Following this procedure, we then examined patterns of clustering in the randomly sampled populations and identified intermediate individuals falling between two genotypic clusters. To further test for the presence of hybrid individuals, we performed two additional ancestry analyses on the sympatric and contact-zone samples (Cherry Burton Road *n* = 20 WGS, Canal Lake *n* = 70 [20 WGS + 50 RAD], Salmon River *n* = 39 RAD; 129 individuals total). The allopatric Blues Cove and McIvor Road Junction populations, which form a single tight cluster in the PCA, were excluded from both analyses.

We first performed a supervised Discriminant Analysis of Principal Components (DAPC; Jombart et al. 2010) using the *adegenet* R package. RAD ecotype labels in our sample sheet were originally assigned post-hoc by PCA, so training a DAPC on those labels would simply rediscover the PCA boundary. To avoid this circularity, we trained the DAPC on the 40 WGS samples only, using their *a priori* morphological labels (Cherry Burton common, Canal Lake white), and then predicted the ecotype of the 89 RAD-sequenced individuals as held-out via predict.dapc(). The number of retained principal components was chosen by cross-validation within the WGS training set (xvalDapc, 100 replicates, 90/10 split, n.pca.max = 30); n.da was fixed at 1 (two-group fit). Mean-imputation of missing genotypes pulls held-out RAD samples toward the discriminant midpoint, producing an apparent excess of intermediate individuals. To minimize this artifact, SNPs were restricted to those with ≤ 5% missingness in the RAD subset prior to imputation (2,318 SNPs after filtering).

We then performed sparse non-negative matrix factorization (sNMF; Frichot et al. 2014) using the *LEA* R package. sNMF estimates individual ancestry proportions without prior labels and is therefore an unsupervised complement to the supervised DAPC. Per-SNP missingness and minor allele frequency were recomputed on the 129-sample mainland subset (biallelic, MAF ≥ 0.03, depth/GQ/missingness/allele-balance filters as above), and no LD pruning was applied. We ran snmf() for *K* = 1–6 with 100 replicates per *K* and chose the best run per *K* by minimum cross-entropy.

### Statistical analysis

We compared the survival of father-reared offspring of F1 hybrid males (♂Cx♀W and ♀Cx♂W) to the survival of father-reared offspring of F0 fathers (common and white). Cross types were lumped to improve statistical power in survival analyses. The survival of father-reared offspring was estimated by physically counting the number of embryos remaining in the tank four days after fertilization, as described above. To confirm that intrinsic hybrid incompatibilities and/or sterility do not contribute to reproductive isolation in this system, we also compared the survival of artificially-reared embryos sired by F0 vs F1 hybrid males. The survival of artificially-reared offspring was quantified by counting the number of surviving embryos after four days of artificial incubation. Differences in clutch size were accounted for statistically by dividing the number of surviving embryos by the original clutch mass in father-reared clutches and by dividing the number of surviving embryos by the original number of eggs in the clutch in artificially-reared clutches. Father-reared offspring survival (surviving embryo count per mg clutch mass) was normally distributed and was analyzed via a linear model. Survival of artificially-reared offspring (surviving embryos per total eggs) was highly right-skewed and fit a beta distribution according to goodness of fit statistics. We therefore analyzed artificially-reared survival as a proportion in a Beta regression model using the *betareg* function of the package *betareg*. A small number (0.001) was added or subtracted to the proportion if necessary to ensure values were between 0 and 1. The paternal cross type (F0, F1), maternal cross type (common, white), and their interaction were included as fixed effects in both models. In both cases, we compared full and reduced models with AIC and likelihood ratio tests and retained models with the best fit.

In the filial cannibalism experiment, the pre-fertilization mass of males was recorded but the mass and egg count of female clutches was unavailable. Filial cannibalism was therefore quantified using raw counts of eggs found in the stomach of a male. This dataset was non-normal and analyzed with non-parametric statistical tests.

Courtship success (number of trials before successfully mating) data was discrete and right-skewed, so differences among male cross types were tested via non-parametric Kruskal-Wallis tests. Separate tests were performed for each female cross type. Nest architecture scores were nonnormal and analyzed with nonparametric tests (Kruskal-Wallis and Spearman rank correlations). Parenting behaviors of stickleback are known to differ across time and populations (Behrens et al. 2024), so we examined two timepoints that are known to differ across groups: immediately after fertilization (day 0) and four days after fertilization (day 4). Parenting behaviors were fitted to linear models with the cross types of the male, female, and their interaction as fixed effects. Dispersal data was overdispersed and zero-inflated but approximated a normal distribution after log+1 transformation. AIC-based model comparison indicated a greater fit with transformed values in a linear model than raw values in a negative binomial model (ΔAIC = 92). Similarly, fanning data was not normally distributed, but square-root transformation yielded significantly stronger support than a Gamma GLM (ΔAIC = 163). Models of transformed data were therefore retained for both behaviors. Full and additive models were also compared for both behaviors. Interaction terms were not supported for dispersal (ΔAIC = 1.994) or day 0 fanning (ΔAIC = 1.744) behaviors so additive models were retained. However, the interaction term improved the fit for day 4 fanning behaviors (ΔAIC = 1.019), so the full model was retained.

Analyses were performed in R version 4.2.1 (R development core, 2011) and visualized with the package ‘ggplot2’ (Wickham 2011).

## RESULTS

### Offspring of F1 hybrid males had low survival when reared by their father

To test the hypothesis that offspring reared by F1 hybrid males suffer greater mortality than offspring reared by F0 males, we compared the survival of father-reared offspring of F1 hybrid males (♂C x ♀W and ♀C x ♂W) to the survival of father-reared offspring that were reared by F0 fathers (common and white). The interaction term was not supported (ΔAIC =1.999), so the additive model was retained. Paternal cross type had a significant effect on survival in father-reared clutches (Fig. 1d; male F_1,33_ = 4.32, p = 0.046; female F_1,33_ = 1.91, p = 0.176), with offspring of F0 fathers exhibiting higher survival than offspring of F1 hybrid fathers.

**Figure 1.**
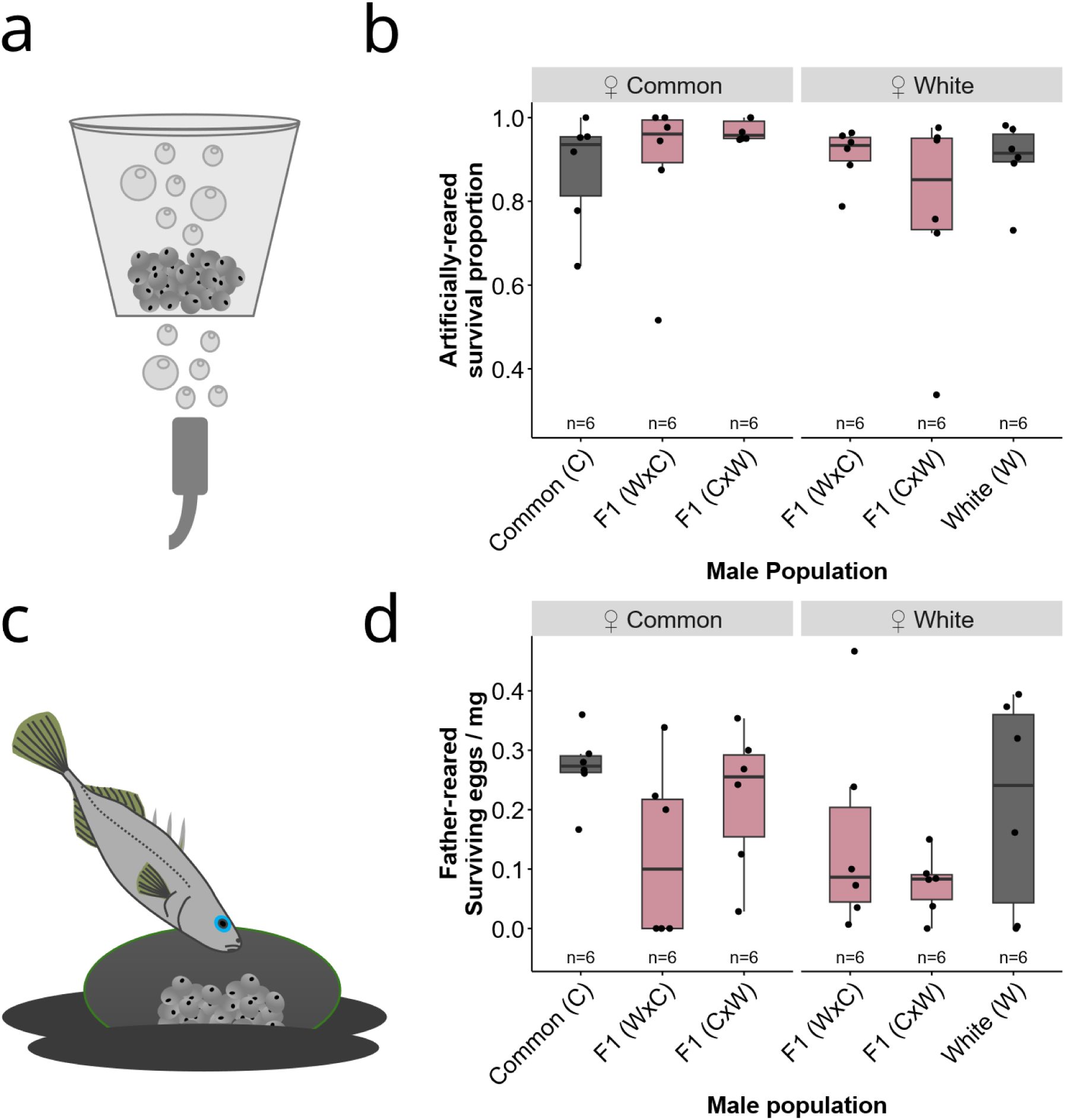
Survival of offspring under artificial and father-rearing. (a) embryos undergo artificial-rearing over a bubbling airstone, and (b) offspring do not differ in survival under artificial-rearing conditions. (c) Father-reared offspring remain in the nest, and (d) model testing suggests that offspring from F0 fathers exhibited significantly higher survival under father rearing conditions. The paternal population is shown on the x-axis and plots are faceted by the maternal population. Boxes are colored by paternal population (F0 in grey, F1 in pink). Each data point represents a clutch, with boxes representing interquartile ranges with the median and whiskers representing 1.5 times interquartile ranges. Post-hoc tests did not detect significant differences among groups.

To confirm that intrinsic hybrid incompatibilities do not contribute to reproductive isolation in this system, we compared the survival of artificially-reared embryos that were sired by F0 versus F1 hybrid males. The interaction term was not supported (ΔAIC =0.753), so the additive model was retained. The survival of artificially-reared embryos differed according to the cross type of the mother but not the father (Fig. 1b; male χ^2^ = 0.09, p = 0.764; female χ^2^ = 4.29, p = 0.038), suggesting a lack of intrinsic hybrid incompatibilities in this system.

Common and white clutches did not significantly differ in mass in artificially-reared (Fig. S1; t = 0.27, df = 32.1, p-value = 0.79) or father-reared clutches (Fig. S1; t_31.9_ = -0.71, p = 0.49), consistent with previous studies (Behrens et al. 2025).

### Dysregulated paternal behavior of F1 hybrid males contributed to low rates of offspring survival

Because offspring reared by F1 hybrid males had lower survival, we next examined the parenting behaviors that may have contributed to this lower survivorship. In general, the parenting behavior of F1 hybrid males was intermediate between whites and commons, and F1 hybrid males exhibited all combinations of elements of the white and common strategies (Fig. 2). For example, some F1 hybrid individuals exhibited relatively high dispersal and fanning rates, while other hybrid individuals exhibited low levels of both behaviors.

**Figure 2.**
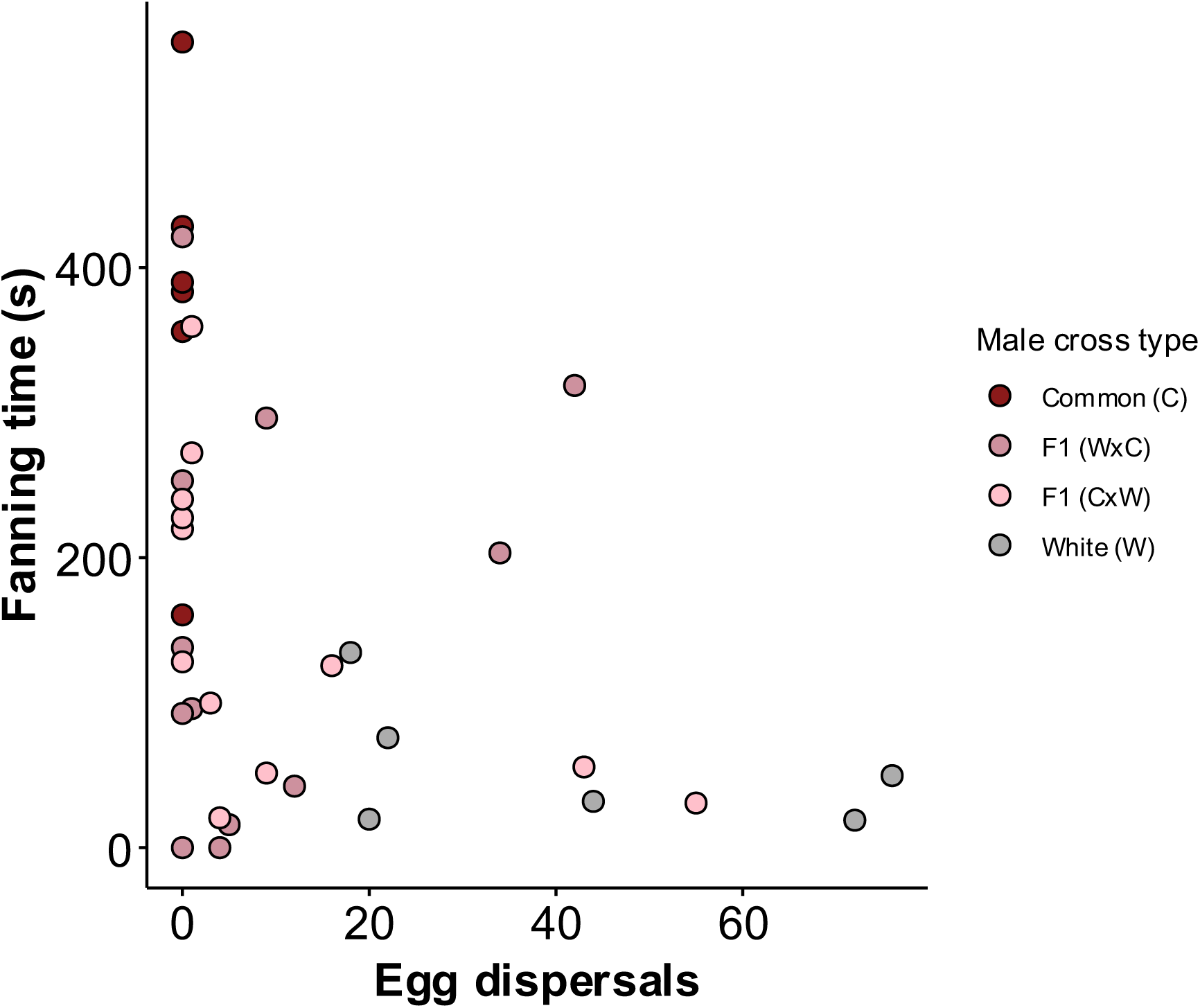
Relationship between the dispersal rate after fertilization and nest fanning four days after fertilization. Data points are colored by cross type. F1 hybrids exhibit a mix of typical white and common behaviors.

Male commons never dispersed embryos (Fig. 3a; Video S1), consistent with previous studies (Blouw 1996; Behrens et al. 2024). In contrast, male whites and F1 hybrids dispersed embryos from their nest (Video S2; Video S3), but the number of dispersals depended on the cross type of both the male and female (Fig. 3a; male F_3,31_ = 15.25, p < 0.001; female F_1,31_ = 25.20, p < 0.001), with embryos from female whites being more readily dispersed than embryos from female commons.

**Figure 3.**
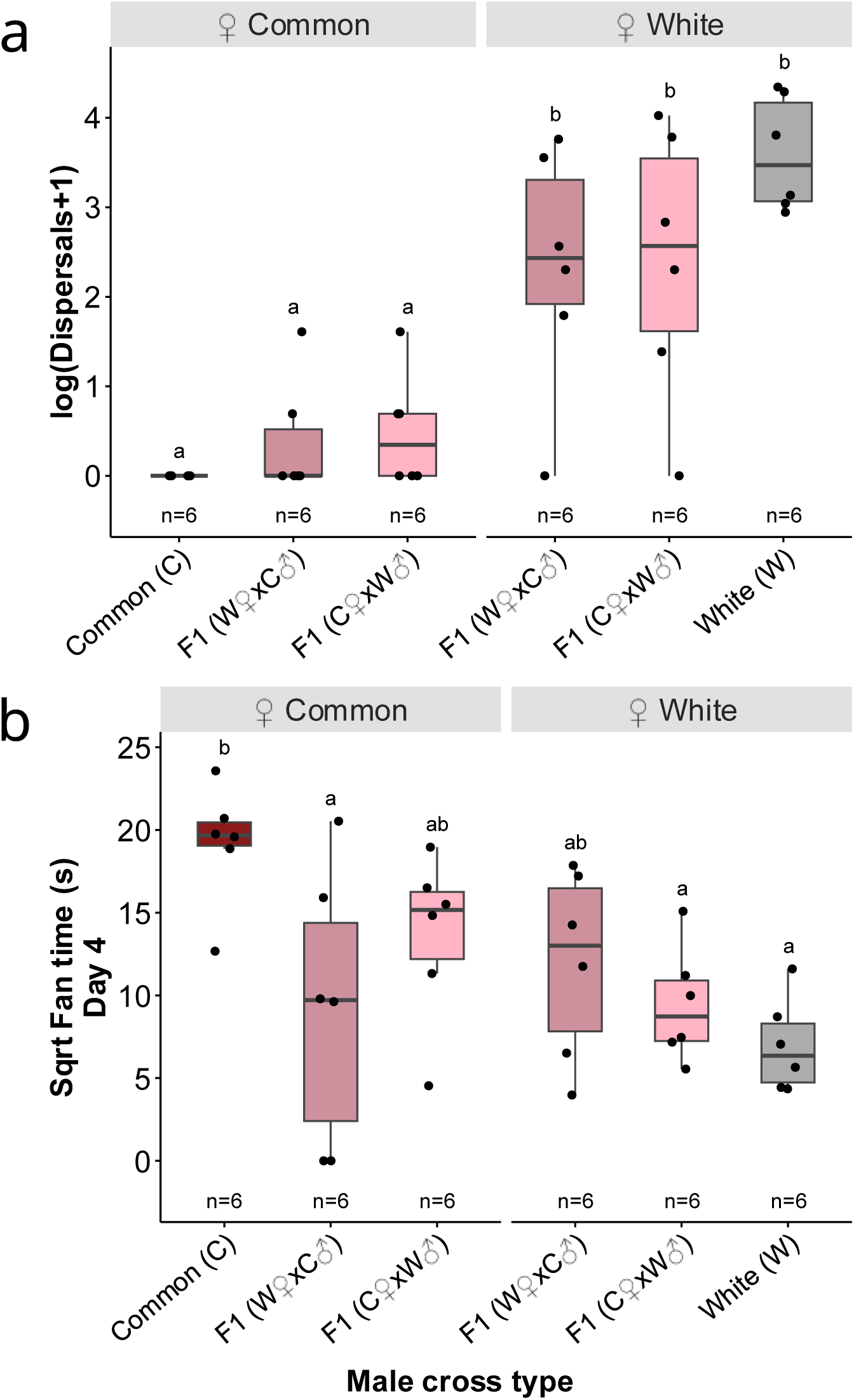
Differences in paternal care across male and female cross types. (a) Fathers dispersed eggs from white clutches more frequently than eggs from common clutches. (b) Male commons fanned their eggs more than male whites, and F1 hybrids were often intermediate. Plots are faceted by female cross type. Each data point represents a male, with boxes representing interquartile ranges with the median and whiskers representing 1.5 times interquartile ranges. Significance lettering reflects Tukey HSD post-hoc tests.

As embryos develop, stickleback fathers typically increase their fanning rate to provide aeration. We did not detect differences in fanning immediately after fertilization (male F_3,31_ = 1.51, p = 0.232; female F_1,31_ = 0.89, p = 0.353) but male commons exhibited significantly higher rates of fanning than male whites and F1 hybrids at four days after fertilization (Fig. 3b; male F_3,30_ = 6.09, p = 0.002; female F_1,30_ = 0.14, p = 0.71; male:female F_1,30_ = 2.62, p = 0.116). Whether the male was mated to a white or common female did not influence levels of fanning behavior in F1 hybrids (Fig. 3b).

One possible explanation for the behavioral sterility of F1 hybrid males is that they failed to modulate their behavior in response to cues from embryos by, for example, failing to fan their embryos, which caused the embryos to die. Indeed, clutches from F1 hybrids that received more fanning had higher rates of survival (dispersal F_1,20_ = 2.93, p = 0.102; fanning F_1,20_ = 11.29, p = 0.003; dispersal:fanning F_1,20_ = 4.60, p = 0.044). However, contrary to this hypothesis, fanning was significantly associated with the number of embryos in a nest (Fig. 4; r = 0.68, p < 0.001), and both F1 hybrid crosses appeared to modulate their fanning rates according to the number of remaining embryos in their nests (F1 WxC r = 0.55, p = 0.062; F1 CxW r = 0.70, p = 0.011). Together, these results suggest that F1 hybrid males are capable of providing care; they attended to cues of embryos in their nest and modulated their behavior accordingly, but their offspring did not survive for other reasons.

**Figure 4.**
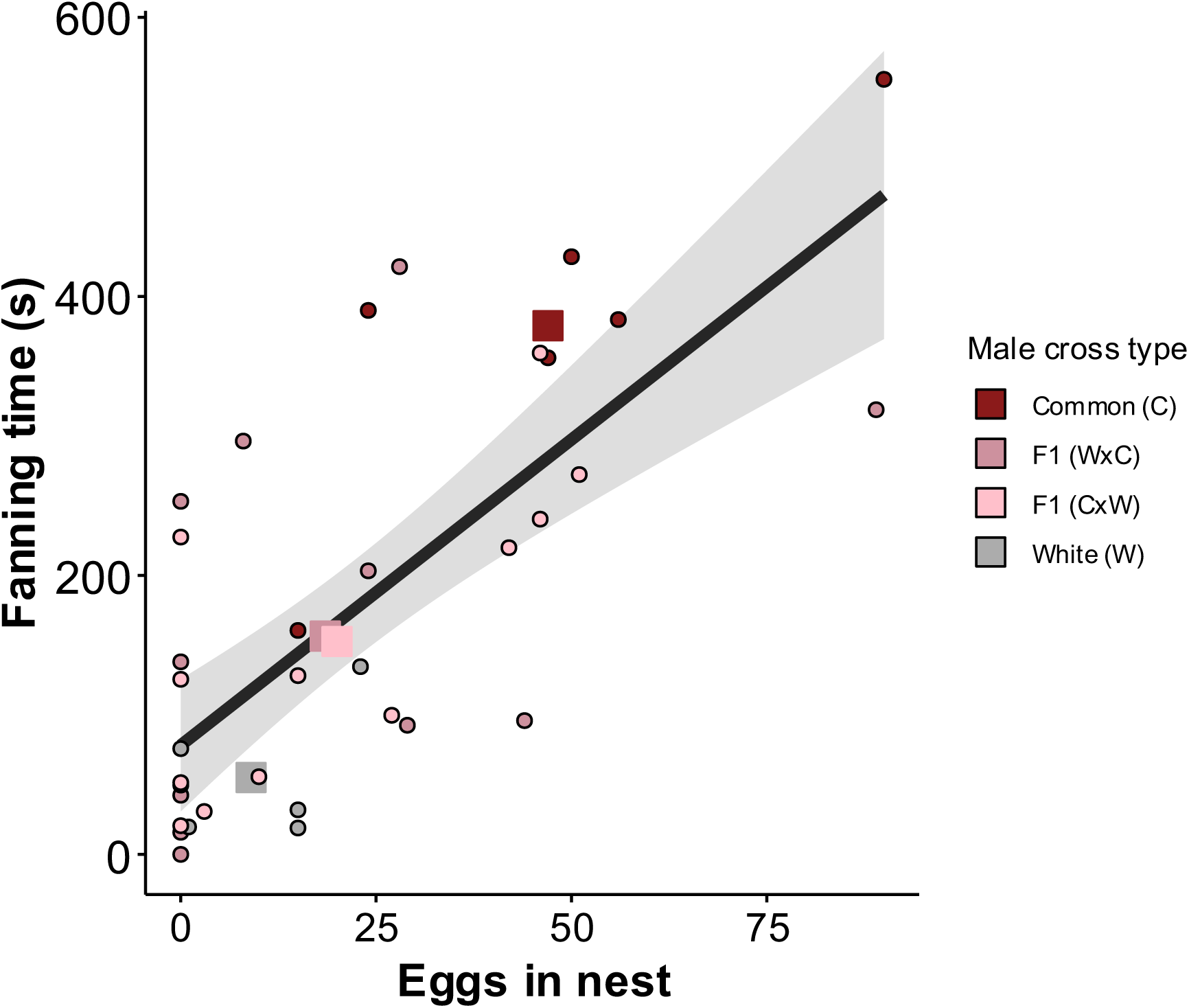
Relationship between fanning rate and the surviving embryos in a nest four days after fertilization. Data points are colored by cross type. Small circles represent individuals and large squares represent group means. The regression line is shown with its 95% confidence interval. Stickleback fathers fan more when more embryos are present in the nest.

Therefore, we investigated another behavioral mechanism that may have contributed to the low survival of offspring of F1 hybrid father-reared offspring: filial cannibalism. Consistent with the hypothesis that hybrid fathers have poor reproductive success due to filial cannibalism, F2 hybrid males had many more eggs/embryos in their stomachs (μ = 26.5, SD 18.3, n=77) relative to either white (μ = 6.77, SD 6.55, n=22) or common (μ = 2.14, SD 3.04, n=22) males (Fig. 5; Kruskal-Wallis, χ^2^ = 58.33, df = 2, p < 0.001). We found no evidence that F2 hybrids engaged in high rates of filial cannibalism because they were in poor condition or small: both male commons and F2 hybrids were heavier than male whites (Kruskal-Wallis, χ^2^ = 6.40, df = 2, p < 0.041), and F2 hybrids consumed significantly more eggs than commons or whites even when corrected for body mass (Kruskal-Wallis, χ^2^ = 30.85, df = 2, p < 0.001). We detected a large proportion of fertilized embryos in F2 males’ stomachs (36% of eggs were unbroken, 67% of unbroken eggs were fertilized). Although the fertilization status of the majority of eggs was unclear due to mastication, the results suggest that hybrid males may not preferentially consume unfertilized eggs that could threaten the entire clutch. Instead, they appear to have consumed eggs regardless of viability.

**Figure 5.**
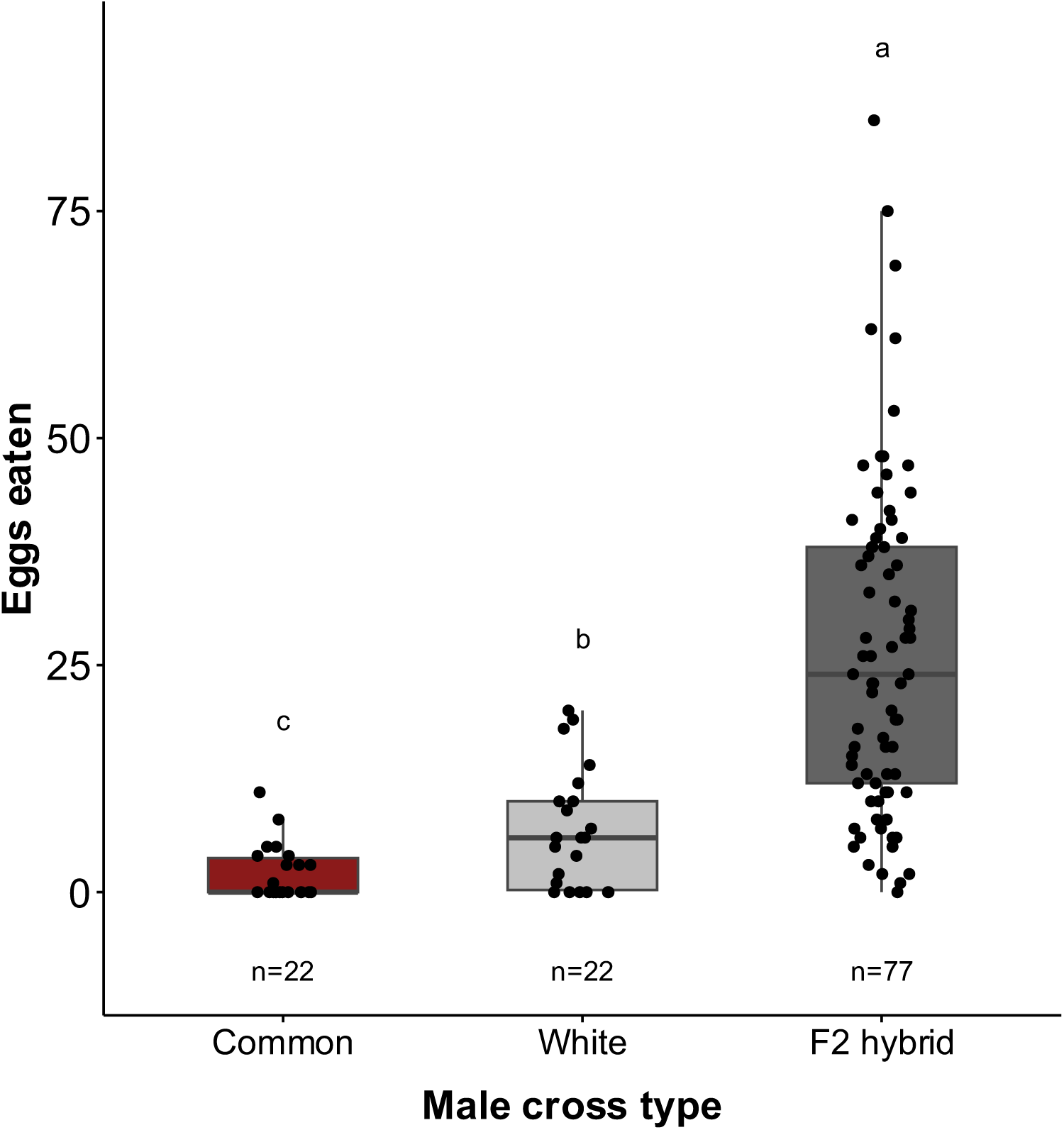
Cannibalism rates in stickleback fathers. Common stickleback consume fewer eggs than white stickleback, and F2 hybrids consume significantly more eggs than either of the F0 cross types. Each data point represents a male, with boxes representing interquartile ranges with the median and whiskers representing 1.5 times interquartile ranges. Significance lettering reflects post-hoc Dunn’s tests.

### Little evidence for mate choice against F1 hybrid males or nest architecture as an isolating barrier

There was no evidence that F1 hybrid males were less attractive to females: mating success (the number of trials before successfully mating) did not differ among male cross types when paired with female commons (Kruskal-Wallis χ^2^ = 0.54, df =2, p=0.760, n=6 per group) or female whites (Kruskal-Wallis χ^2^ =2.40, df=2, p=0.302, n=6 per group), suggesting that male F1 hybrids can readily mate with F0 females.

Since divergent nest architecture could act as a prezygotic isolating barrier if female whites and commons exhibit different nest preferences, we investigated if there was any evidence for assortative mating based on nest architecture. Consistent with previous studies (Behrens et al. 2024), nest architecture significantly differed according to male cross type (Fig. S2; Kruskal-Wallis χ^2^ = 10.55, df = 3, p=0.014). Whites built high-scoring nests (tall and loose), commons build low-scoring nests (compact and sandy), and F1 hybrids built intermediate nests. There was no evidence that the nest score of F1 hybrid males influenced mating success with female commons (*ρ* = -0.14, p=0.726, n=9) or whites (*ρ* = -0.28, p=0.407, n=11).

Nest architecture might also be important as a postzygotic isolating barrier if, for example, it is easier for males to disperse offspring from white-like nests, or to fan embryos in common-like nests. We found no evidence that F1 hybrid nest score was correlated with rates of dispersal when mated to female whites (*ρ* = -0.48, p=0.135, n=11). However, F1 hybrid nest score was significantly associated with rates of fanning when mated to female commons (*ρ* = - 0.85, p=0.004, n=9), with common-like nests receiving greater rates of fanning.

### Whites and commons hybridize in the wild

Despite significant phenotypic divergence, white and common stickleback exhibit low genetic divergence and are likely experiencing ongoing gene flow (Samuk 2016). If postzygotic behavioral isolation via dysregulated paternal care is acting as a behavioral isolating mechanism, then F1 hybrids should be detectable in the wild and behavioral sterility could provide an explanation for the limited gene flow among populations. Consistent with this line of reasoning, while wild-caught commons and whites clustered into distinct genetic groups, putative hybrids were detected as genetically intermediate individuals (Figure 6). The presence of these individuals is the first genetic evidence of ongoing, natural hybridization in this system, and suggests that common and white stickleback naturally hybridize in the wild at low levels, with 2.5% of randomly sampled individuals classified as putative hybrids across sympatric sites (Fig. 6). Hybrid class (e.g. F1, F2, backcross, etc.) was unable to be determined due to low genetic differentiation between commons and whites. Randomly sampled individuals at sites known to contain only common sticklebacks displayed a single genotypic cluster (Fig. S3; BC and MJ clusters), suggesting that the clustering and intermediates found at the white/common sympatric sites are not an artifact of sampling.

**Figure 6.**
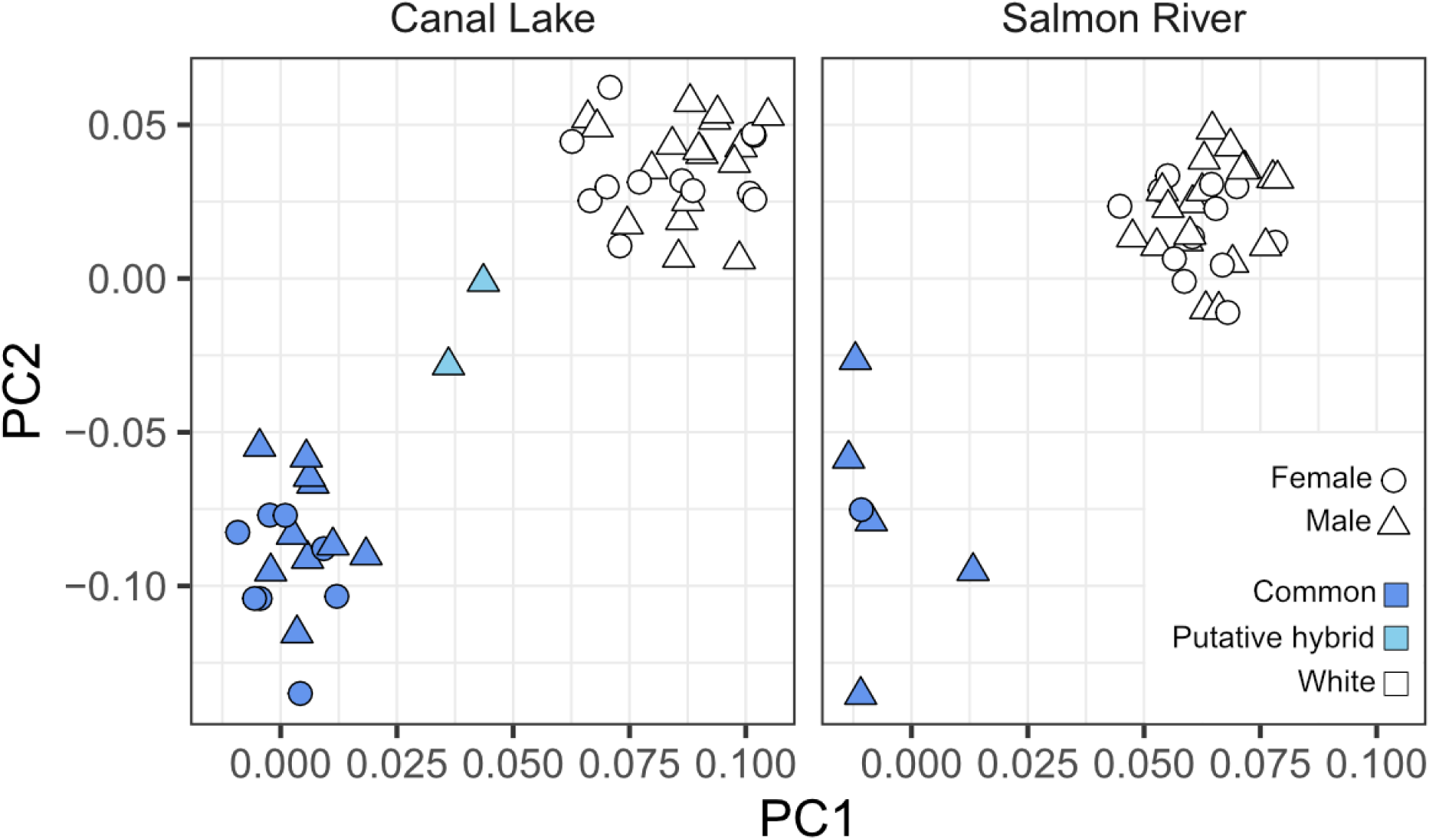
A principal component analysis of RADseq-derived SNPs from randomly sampled stickleback individuals in two locations where white and common sticklebacks are sympatric (Canal Lake and Salmon River, Nova Scotia). Points are colored by ecotype (white, common) and sex (male, female) at each site, with putative hybrids indicated. Points are plotted in the same latent space as Figure S3.

We further tested for the presence of hybrid individuals using two additional ancestry analyses on the sympatric and contact-zone samples (Canal Lake and Salmon River, with Cherry Burton Road as a common-only reference; 129 individuals total). The first was a supervised Discriminant Analysis of Principal Components (DAPC; Jombart et al. 2010), trained on the 40 WGS samples using their *a priori* morphological ecotype labels and then used to predict the ecotype of the 89 RAD-sequenced individuals as held-out. The second was a sparse non-negative matrix factorization (sNMF; Frichot et al. 2014), an unsupervised model-based estimator of individual ancestry proportions that requires no prior labels. Both analyses recovered the same axis of common/white differentiation as the PCA, and both identified genetically intermediate individuals concentrated at Salmon River and among the RAD-sampled fish from Canal Lake (Fig. S4, Fig. S5). All individuals identified as putative hybrids by the PCA were also identified as intermediate by both methods, and the direction of admixture (common-leaning vs. white-leaning) agreed across analyses. sNMF identified several further admixed individuals that the PCA did not resolve, suggesting that the ∼2.5% figure reported above is a conservative lower bound on the rate of hybridization in these sympatric populations.

## DISCUSSION

Speciation is one of the most vexing problems in evolutionary biology, as it is critical for understanding the origins and maintenance of biodiversity but is challenging to study. An essential goal of the field is to identify how traits contribute to speciation, and the study of reproductive isolating mechanisms in the early stages of divergence between populations (before confounding differences have accumulated) has helped to advance our understanding of the speciation process. The radiation of three-spined stickleback has been an important model in speciation research for these reasons (McKinnon and Rundle 2002).

This study leverages the white-common stickleback system to present empirical evidence for a novel behavioral postzygotic reproductive isolating mechanism: behavioral sterility via dysregulated paternal care. The possibility that disrupted parental behavior might contribute to reproductive isolation was first suggested by Buckley (1966), who found that female hybrid *Agapornis* parrots display altered nest-building and, if their eggs were fertilized, poor parental care that led to clutch failure. However, these hybrid parrots also suffered almost complete physiological sterility and poor viability (Buckley 1969), and therefore their altered parental behavior likely does little to prevent gene flow. In contrast, in the white-common stickleback system, there is no evidence for intrinsic postzygotic barriers like hybrid gametic sterility or inviability. Behavioral sterility in the form of dysregulated paternal behavior in hybrid male stickleback could, therefore, play an important role as a reproductive isolating barrier in this species, and these results add empirical support to theoretical models that have suggested such a pattern (Reyes et al. 2025).

Theory often predicts that isolating mechanisms acting relatively early in reproduction (e.g., timing of breeding, mate choice) are more effective isolating barriers than mechanisms that act later (e.g. hybrid genetic incompatibilities; Mayr 1963; Coyne and Orr 2004), but models suggest that partial prezygotic barriers, like assortative mating, can be ineffective without accompanying postzygotic isolation (Irwin 2020). In the white-common stickleback system, prezygotic barriers appear to be minimal and/or evidence for them is inconclusive (Blouw and Hagen 1990; Corney and Weir 2023; Behrens et al. 2024), therefore behavioral sterility via dysregulated paternal care may contribute to reproductive isolation in this system, and this (in addition to very recent divergence) could help explain why whites and commons are only weakly genetically differentiated. However, the low rate of hybridization in the wild (∼2.5%) suggests that prezygotic barriers, microhabitat preferences, and sexual selection against hybrids may be stronger in natural environments, as is observed in other stickleback systems (Vamosi and Schluter 1999; Bolnick et al. 2015). For example, variation in nest architecture may act as a prezygotic barrier (potential mates prefer specific designs), and as a postzygotic barrier (common-like nests received greater rates of fanning). Location-specific factors (e.g. habitat variation, population demographics, demographic history) may also influence hybridization rates, as hybrids were only detected in one of the two sympatric sites, but increased sampling would be required to test this, Additionally, sexually imprinted traits are known to contribute to speciation in stickleback (Kozak et al. 2011), meaning daughters imprinted on a father’s traits may affect mate preference and further facilitate speciation.

The specific genetic and behavioral causes of dysregulated paternal behavior in F1 hybrids are not known, but we suspect that a primary causes of mortality of offspring reared by hybrid males are pathogens, caused by the father’s failure to perform an appropriate combination of behaviors to combat fungus, and/or filial cannibalism. To prevent fungal infection, male commons tend the nest and remove dead and decaying eggs. In contrast, male whites prevent the spread of pathogens by breaking apart the clutch and scattering the embryos in small clumps. Hybrid fathers performed a mix of behaviors that typify the white and common strategies, and it is possible that these low/combinatorial levels of behavior may have directly contributed to the death of offspring due to fungal infection. Indeed, dysregulated parental care may have stronger effects in natural conditions, where pathogens and predators are ubiquitous, and field studies could further clarify the strength of postzygotic isolation in this system.

While the care provided by hybrid fathers may have contributed to the low survival of offspring, we also provide evidence suggesting that white and common reproductive strategies may be incompatible at the genetic level, causing high rates of filial cannibalism in F2 hybrids. Filial cannibalism could also explain why levels of paternal behavior by F1 hybrid fathers were often low relative to white or common fathers: embryos elicit caregiving behaviors (Fig 4, see also van Iersel 1953), and if the F1 hybrid fathers consumed embryos, then there were fewer embryos to fan. Note that although significant cannibalism was observed by F2 hybrids, cannibalism by F1 hybrid fathers was not quantified in this study; clarifying the link between cannibalism rate and dysregulated parental care is a priority for future studies.

The high rates of cannibalism by hybrid fathers could reflect dysregulation of overlapping neural circuits involved in feeding, infant-directed aggression and parenting. Growing evidence in vertebrates suggests that core conserved neuropeptides, such as galanin, mediate both feeding and care (O’Rourke and Renn 2015; Fischer and O’Connell 2017). For example, optogenetic stimulation of preoptic galanin neurons was sufficient to switch the behavior of laboratory male mice from infanticidal to active parental behavior (Wu et al. 2014). Intriguingly, galanin is also differentially expressed during paternal care (Bukhari et al. 2019) and preoptic galanin neurons are differentially activated during caregiving in commons compared to whites (Maciejewski et al. 2025). Therefore, we hypothesize that genetic incompatibilities may have disrupted parental and feeding circuits, causing hybrid stickleback fathers to consume rather than care for their embryos. However, hybrid filial cannibalism was only quantified across a single F2 hybrid mapping family, and it is possible that genetically-based deleterious effects were amplified or diminished in our dataset. Additional studies are required to better understand how neuronal circuits and genomic loci contribute to cannibalism in this system, especially in wild populations.

Consistent with the hypothesis that hybrid males engaged in higher than typical rates of filial cannibalism due to disrupted neural circuits, hybrid male fathers did not preferentially consume unhealthy embryos or unfertilized eggs that were dead or decaying. Instead, many of the embryos found in hybrid males’ stomachs were fertilized and developing normally, and filial cannibalism was quantified within one hour of mating, which is too soon for fungal infections to start or spread. Indeed, under natural circumstances, rates of filial cannibalism by hybrid fathers may be even higher because their embryos fail to develop due to altered care, per above, or because their spiggin – a glue that stickleback fathers use to hold the nest together – might lack antimicrobial properties (Little et al. 2008).

This study focuses on male parental behavior, but females also likely play a critical role in driving reproductive isolation in this system. Typically, female stickleback produce viscous ovarian fluid that holds eggs together within a clutch. In contrast, the ovarian fluid of female whites is relatively thin (Grant 1993), which may facilitate the egg scattering behavior of male whites after fertilization and prevents the offspring from staying “all in the same basket”, and thereby vulnerable to total loss by predation or fungus. In support of this hypothesis, our results suggest that the eggs of female whites were easier for males to disperse. These findings prompt future investigation into the adhesiveness of F1 hybrid female clutches (preliminary studies suggest they are intermediate; Bagazinski, A., Maciejewski, M.F., Bell, A.M., personal communication) and raise interesting questions about the outcome of white x common matings and the direction of introgression. For example, if hybridization is limited by hybrid males, but not hybrid females, then females could be driving gene flow between populations. Future studies should examine the reproductive biology of hybrid females (mate choice behavior, embryo survival, clutch viscosity, etc.) to investigate this possibility.

More than 150 years after Darwin’s pivotal discoveries, speciation is still challenging to understand. Identifying the processes involved in speciation is difficult, in part because it requires careful examination of natural history to understand the biology of the study system, and because speciation research frequently requires a retrospective view of events that occurred thousands or millions of years in the past. Identifying reproductive isolating mechanisms and quantifying their effects in systems is therefore key to understanding the speciation process. Our results contribute to this field by suggesting that dysregulated parental care can act as an underappreciated reproductive isolating barrier in stickleback and other animals.

## Author contributions

C.B. and A.M.B conceptualized the project. A.S., G.N.L., and K.S. performed field collections in 2023 and reared organisms. G.N.L. and K.S. performed field collections (RADseq) in 2022. C.B. collected behavioral data. C.B. and M.F.M. collected cannibalism data. G.N.L. performed DNA extractions and normalizations. G.N.L. and G.L.-R. performed RAD-seq library preps. A.S and K.S processed the sequencing data for population genetic analyses. C.B. and K.S. performed analyses and data visualization. C.B. and A.M.B wrote the original draft. C.B., K.S., and A.M.B reviewed and edited the manuscript.

## Data Availability Statement

Raw RAD sequence data are deposited in the NCBI Sequence Read Archive under BioProject: PRJNA1244444. Raw WGS sequence data are deposited under BioProject: PRJNA1244922. Accession numbers for each sample are listed in Table S1. Behavioral data and code will be made available at Dryad.

## Acknowledgements

We thank members of the Bell lab and anonymous reviewers for insightful comments that improved the manuscript, and we are grateful to Mark Hauber for advice on designing the experiment. This work was supported by the National Institute of General Medical Sciences of the National Institutes of Health under awards 1R35GM139597 and 2R01GM082937-06A1, and by a National Science Foundation Graduate Research Fellowship (DGE-1326120) to G.L.-R. Computations were performed using the computer clusters and data storage resources of the HPCC, which were funded by grants from NSF (MRI-2215705, MRI-1429826) and NIH (1S10OD016290-01A1).

## Notes

### Competing Interest Statement

The authors have declared no competing interest.

### Summary of Updates

Additional ancestry analyses (DAPC, sNMF) have been performed and added in support of the original PCA-based analyses. This is reflected in new text and supplemental figures. The introduction and discussion have been edited for clarity. Methods have also been expanded to describe genomic and behavioral analyses in greater detail.

