## Supplement for "Evidence of parental care as a newly identified reproductive isolating barrier"

**Supplemental**

**Method S1:** Animal husbandry

Fry were reared in 5-gallon aquariums and later moved to 20-gallon aquariums at the University of California Riverside (University of California Riverside IACUC #83). Fish were maintained at 17°C with a 12:12 (L:D) photoperiod and fed brine shrimp, blood worms, and mysis shrimp daily. In the spring of 2024, fish were transported to facilities at the University of Illinois Urbana-Champaign. Fish were maintained at summer conditions (20°C with 16:8 (L:D) photoperiod) and fed a mixture of blood worms, mysis shrimp, brine shrimp, and cyclops daily. Experimental protocols were approved by the University of Illinois Urbana-Champaign IACUC Protocol #24049.

**Method S2:** Sample collection

Stickleback were trapped using unbaited ½" mesh Gee Minnow traps Twelve minnow traps were set at each site for 8-12 hours, after which their contents were placed into aerated buckets containing fresh sea water. All non-three-spined individuals were removed from the buckets and returned to the trapping sites. Fifty individual three-spined sticklebacks were then randomly removed from the buckets and fin clips of approximately 0.5 cm squared were taken using dissecting scissors. The fin clips were immediately placed in 1.5mL microcentrifuge tubes containing 95% ethanol. All fish were then returned to the trapping sites.

**Method S3:** Variant Calling and Filtering

RADseq and WGS reads were processed independently but followed similar bioinformatics steps unless otherwise stated. Raw RADseq reads were demultiplexed using the process_radtags module in Stacks v 2.6 (Catchen et al. 2013). For both RADseq and WGS datasets, sequence reads were aligned to the three-spined stickleback reference genome v5 (Nath et al. 2021) using bwa mem (Li and Durbin 2009). Duplicate reads in the WGS data were removed using Picard (2019). Variant calling was performed using GATK HaplotypeCaller (McKenna et al. 2010). All prior steps were carried out using the snakemake pipeline, grenepipe (Czech and Exposito-Alonso 2022). The resulting GVCFs were then merged using the GenomicsDBImport argument in GATK. This was done for each chromosome prior to joint genotyping using GenotypeGVCFs. The resulting vcfs were then subset and filtered in vcftools v 0.1.16 (Danecek et al. 2011). In general, we retained variants that were biallelic snps and followed the GATK best practices hard filters. The separate RADseq and WGS vcfs were then intersected to identify regions shared between both datasets using bedtools v 2.31.0 intersect (Quinlan and Hall 2010). The intersected VCFs containing shared sites were then merged together using bcftools v 1.16 (Li 2011). Finally, we filtered the intersected VCF using snpfiltR (DeRaad 2023) with the following parameters: retaining biallelic SNPs, minimum genotype depth of 5 and quality of 20, maximum missing data 0.85 (i.e. 15% missing data allowed), maximum per sample depth of 40, minimum minor allele count of 1. After intersection and filtering, 4507 high quality SNPs were retained. All variant calling and filtering for SNPs were done on the computer cluster provided by the University of California, Riverside High-Performance Computing Center.

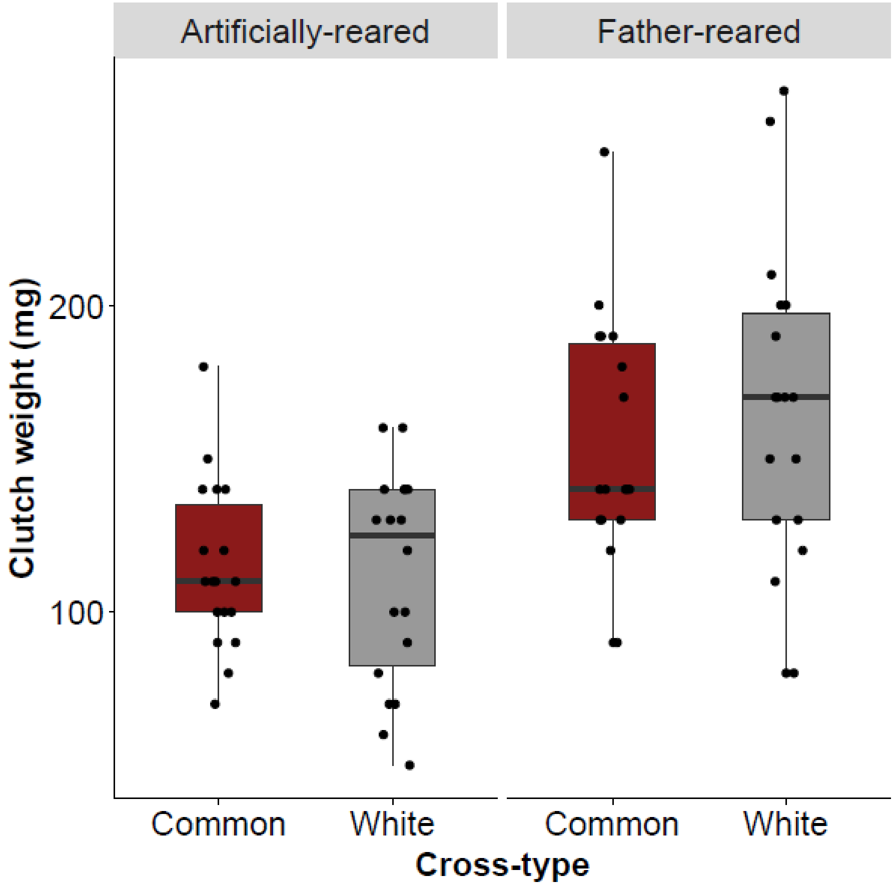


**Figure S1.** Clutch weights from F0 females. Common and white stickleback did not differ in clutch mass, though artificially-fertilized clutches were smaller than father-reared clutches ($t_{60.6}$ = -4.90, p < 0.001). Each data point represents the clutch from an individual female.

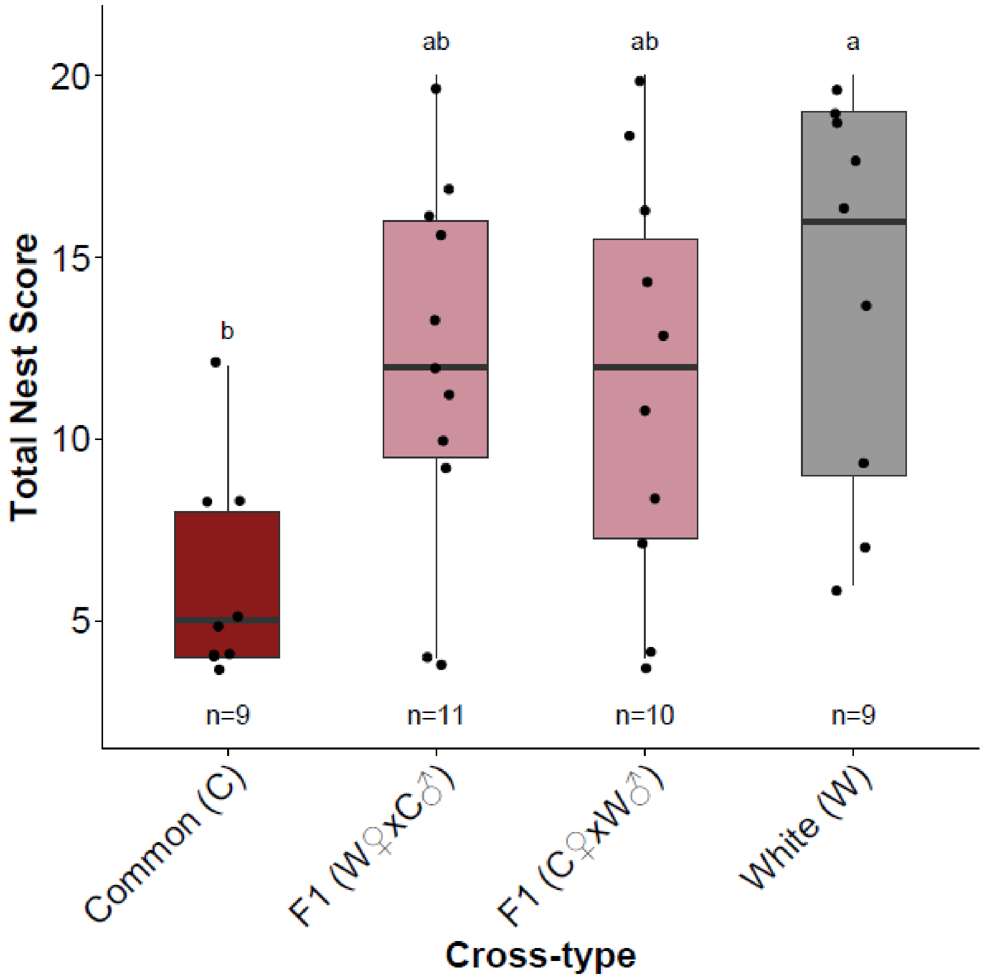


**Figure S2.** Differences in nest architecture. The nest architecture score is a scale (ranging 4-20) incorporating the nest height, nest location, sand content, and nest opening. Higher scores represent tall nests with horizontal openings and no sand. Low scores represent compact nests with vertical openings and large amounts of sand. Each data point represents a nest from a single male. Nests from commons and whites significantly differ in overall architecture, while F1 hybrids are intermediate. Significance lettering reflects Tukey HSD post-hoc tests.

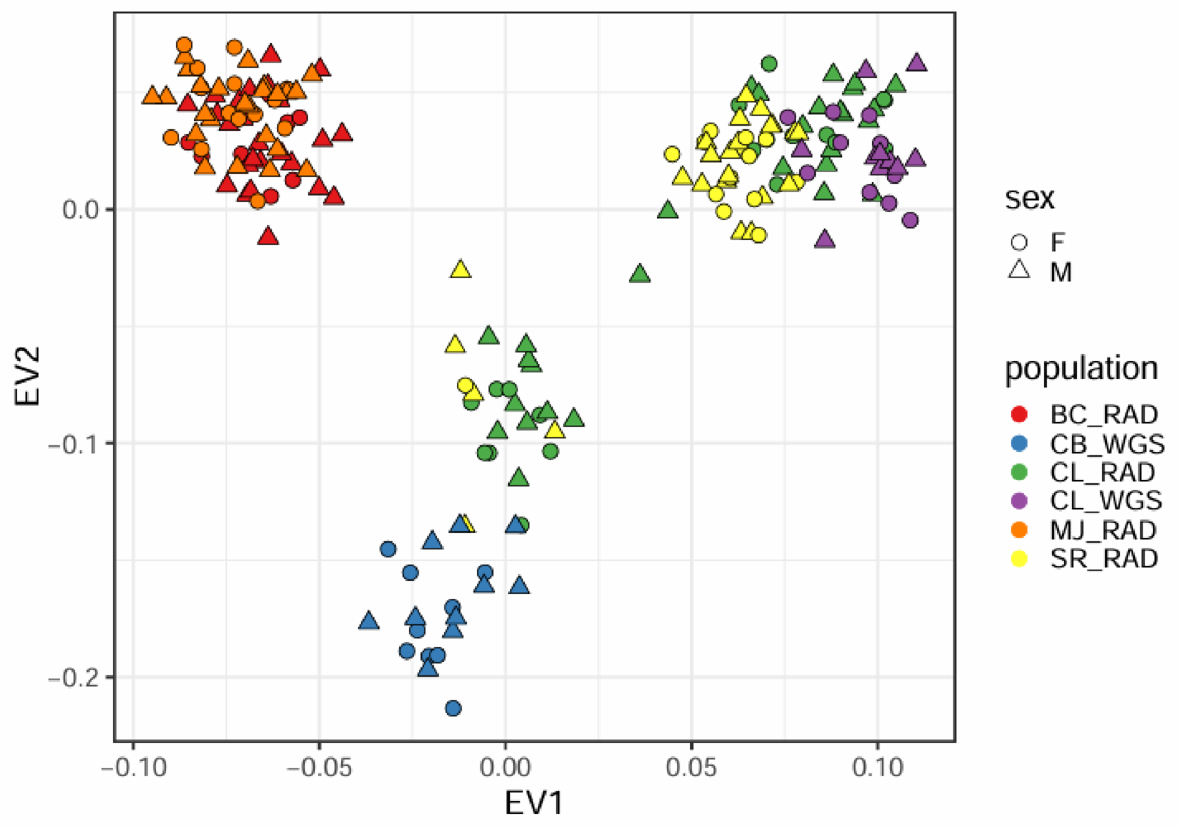


**Figure S3**. Principal component analysis of the jointly filtered SNPs from the randomly sample RADseq individuals ("RAD") and WGS samples with known sex and species identities ("WGS"). Points are colored by population/sequencing strategy, with shape indicating sex. The separate clustering of BC and MJ is due to known geographic structure in the common stickleback population. BC and MJ are both located in the Bras d'Or Lake, an inland sea on Cape Breton Island, whereas CB/CL/SR are found on the mainland of Nova Scotia.

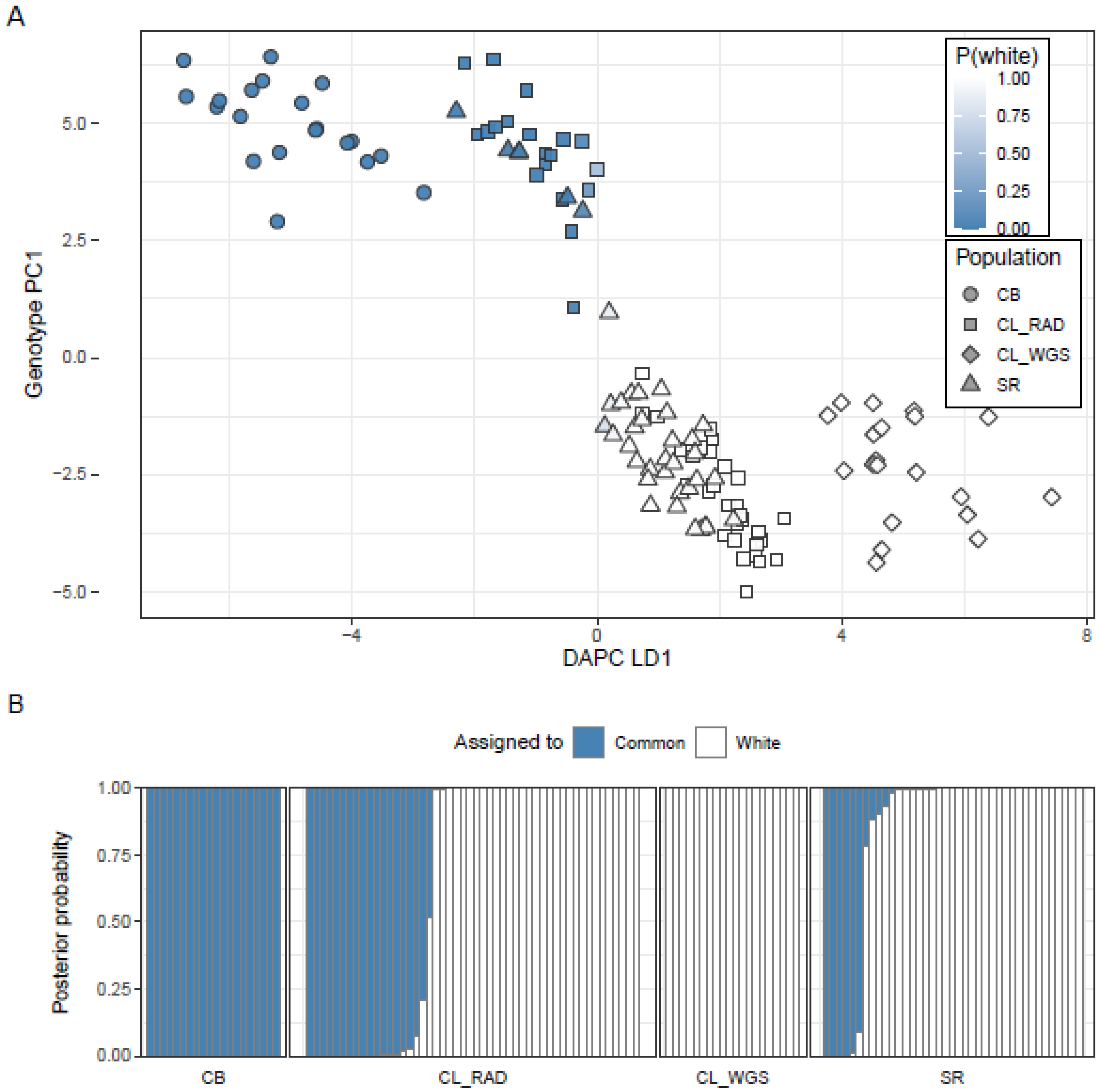


**Figure S4.** Discriminant Analysis of Principal Components (DAPC) of the 129 mainland samples (Cherry Burton Road common WGS [CB, *n* = 20], Canal Lake white WGS [CL_WGS, *n* = 20], Canal Lake RAD [CL_RAD, *n* = 50], Salmon River RAD [SR, *n* = 39]). The DAPC was trained on the 40 WGS samples (CB and CL_WGS) using their *a priori* morphological ecotype labels; the 89 RAD-sequenced individuals (CL_RAD, SR) were then assigned as held-out via predict.dapc(). **(A)** DAPC discriminant score (LD1) plotted against the first principal component of the same genotype matrix. Point shape gives population and point fill gives the posterior probability of assignment to the white ecotype, P(white). The two WGS training populations sit at opposite ends of LD1, and several held-out RAD individuals from CL_RAD and SR have intermediate LD1 scores. **(B)** Per-individual posterior probability of assignment to the common (blue) or white (unfilled) ecotype, sorted within population by P(common). Intermediate posteriors (mixed shading) are present in CL_RAD and SR but absent from the training populations CB and CL_WGS.

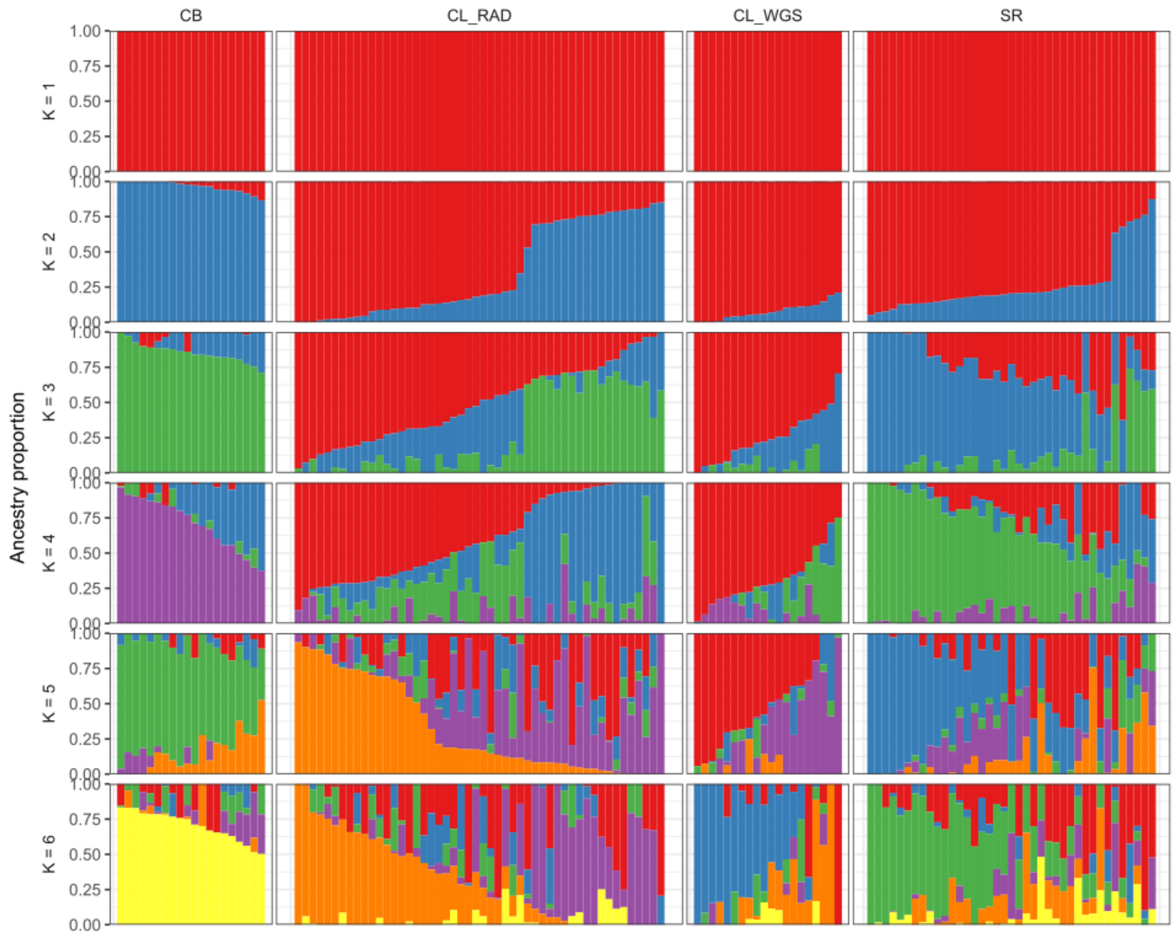


**Figure S5**. Unsupervised ancestry estimates from sparse non-negative matrix factorization (sNMF; LEA::snmf) for the 129 mainland samples (Cherry Burton Road common WGS [CB, $n = 20$], Canal Lake white WGS [CL_WGS, $n = 20$], Canal Lake RAD [CL_RAD, $n = 50$], Salmon River RAD [SR, $n = 39$]). Rows are ancestry proportions for the best run (lowest cross-entropy across 100 replicates) at $K = 1$ to $K = 6$; columns are populations. Each vertical bar is one individual, sorted within population by ancestry proportion at the corresponding $K$. Cross-entropy selects $K = 1$, consistent with weak genetic differentiation between the two ecotypes. At $K = 2$ the model still recovers a clear common/white axis: CB sits almost entirely in one component (median $q = 0.97$) and CL_WGS in the other (median $q = 0.06$), while CL_RAD and SR contain many individuals with intermediate ancestry. These intermediates are the same individuals flagged as putative hybrids by the main-text PCA (Fig. 6) and by the DAPC (Fig. S4).

**Video S1.** A male common stickleback exhibits typical parental care behaviors, including nest attendance, nest fanning, and nest poking. The nest is on the left side of the sandbox.

**Video S2.** A male white stickleback exhibits dispersal behavior. Shortly after fertilizing a clutch, the male removes embryos from his nest and disperses them into the surrounding environment.

**Video S3.** A male F1 hybrid disperses a clutch from a common female. The clutch is sticky and difficult to separate, so the majority of the clutch is removed in a single dispersal. The male then returns to fan his primarily empty nest.
